# DNA methylation clocks for clawed frogs reveal evolutionary conservation of epigenetic ageing

**DOI:** 10.1101/2022.08.02.502561

**Authors:** Joseph A. Zoller, Eleftheria Parasyraki, Ake T. Lu, Amin Haghani, Christof Niehrs, Steve Horvath

## Abstract

DNA methylation-based biomarkers of ageing (epigenetic clocks) have been developed for many mammals, but not yet for amphibian species. We generated DNA methylation data from African clawed frogs (*Xenopus laevis*) and Western clawed frogs (*Xenopus tropicalis*), from adult tissues, whole embryos, and tadpoles. We used an array platform designed for CpGs that are highly conserved in mammals to build multiple DNA methylation-based estimators of age for *Xenopus*. We found that dual species clock could be developed that apply to both humans and frogs (human-clawed frog clocks), and whose high accuracy supports that epigenetic ageing processes are evolutionary conserved outside mammals. Crossing vast evolutionary distances, we characterize age-related CpGs in *Xenopus* and mammalian species. Highly conserved positively age-related CpGs are located in neural-developmental genes such as *uncx, tfap2d* as well as *nr4a2* implicated in age-associated disease. As in human clocks, positively age-related CpGs are associated with Polycomb repressive complex 2 (PRC2) target sites. Negative age-related CpGs are associated with genes involved in synaptic transmission. We conclude that signatures of epigenetic ageing are evolutionary conserved between frogs and mammals and that the associated genes relate to neural-developmental processes, altogether opening opportunities to employ *Xenopus* as a model organism to study development and ageing.

## INTRODUCTION

Stereotypical changes of DNA methylation (DNAm) of cytosine residues within CpG dinucleotides (5-methylcytosine) have emerged as one of the most reliable biomarkers of chronological age in mammals. Indeed, it was long observed that DNAm changes with age in humans and many mammalian species [1–3]. Methylation levels of multiple CpGs can be consolidated to develop highly accurate multivariate age-estimators (epigenetic clocks) for all human tissues [4]. Epigenetic age, as measured by an epigenetic clock, is arguably the most accurate estimator of age in numerous mammalian species including humans [5–8].

Epigenetic clocks for humans have found many biomedical applications, including the measure of age in human clinical trials [7, 9]. This instigated development of similar clocks for mammals such as mice [5, 10–14]. Our Mammalian Methylation Consortium generated data from n=348 mammalian species using the mammalian methylation array platform, which focuses on highly conserved CpGs [15–17]. These methylation data allowed us to characterize age related changes in cytosine methylation levels and to construct multivariate age estimators (referred to as epigenetic clocks) that apply to individual mammalian species. The focus on highly conserved cytosines facilitated the construction of third generation epigenetic clocks that apply to multiple species at the same time [18–23]. This work culminated in the development of pan-mammalian clocks that apply to all mammalian species [16].

The Mammalian Methylation Consortium focused exclusively on mammals. However, DNAme shows age-related signatures also in non-mammalian organisms, including chicken [24], frog [25], and the planktonic crustacean Daphnia [26], raising the question if epigenetic ageing is conserved in non-mammalian vertebrates. Amphibians are a widely used model system because of its experimental tractability and relatively closer evolutionary relationship with humans compared to alternatives such as fish or invertebrates. Among amphibian species, African clawed frog (*Xenopus laevis*) and the Western clawed frog (*Xenopus tropicalis*), are the molecularly best studied. Decades of research have been organized and cataloged in Xenbase [27] and the *X.laevis* and *X.tropicalis* genomes have both been sequenced [28, 29]. *Xenopus* features epigenetic cytosine methylation via DNA methyltransferases *dnmt1* and *dnmt3a* and demethylation by *tet2* and *tet3* demethylases much like mammals [30–35]. As in mammals, *Xenopus* DNAme is correlated with epigenetic histone modifications [35–37] and shows the characteristic promoter hypomethylation [38].

While numerous studies concern the embryonic and larval development of *Xenopus*, the biology of adult ageing has been little analyzed [39–43]. The main difficulty in studying ageing in *Xenopus* is the long lifespan of *X. tropicalis* and *X. laevis*, estimated to be at least 16 and 30 years in captivity, respectively (own experience and [44]). In other amphibian species, studies on adult ageing are mostly limited to lifespan analyses [45–49]. Hence, a reliable epigenetic clock biomarker would benefit the analysis of biological ageing in this important model organism to facilitate DNAm-based assessments upon environmental exposures, for understanding of ageing mechanisms, and for preclinical studies of anti-ageing therapies.

Therefore, in the current article, we focused on addressing the following three questions. First, does the mammalian methylation array platform lend itself for generating informative methylation data in clawed frogs? Second, can one build dual species epigenetic clocks that apply to both clawed frogs and humans at the same time? Third, are age-related DNAme signatures evolutionary conserved beyond mammals? We demonstrate that the answer to all questions turns out to be affirmative. The availability of evolutionary conserved epigenetic clocks (and hence underlying mechanisms), in combination with the experimental tractability of *Xenopus* holds promises to study environmental-as well as metabolic effects on organismic ageing, to gain insights into the underlying mechanisms, and to develop therapeutics.

## METHODS

### Ethics Statement

*Xenopus* experiments were approved by the state review board of Rheinland Pfalz, Germany (Landesuntersuchungsamt, reference number 23177–07/A17-5-002 HP) and performed according to federal and institutional guidelines.

### Frog tissue samples

Adult frogs were obtained from three sources: NASCO Education, European *Xenopus* Resource Centre (EXRC) and National *Xenopus* Resource (NXR). Embryos, tadpoles and juvenile animals were prepared at the Institute of Molecular Biology (IMB) facility in Mainz by *in vitro* fertilization as described in [50]. *X. laevis* and *X.tropicalis* embryos were cultivated in 0.1x Barth’s solution. *Xenopus laevis* were kept at 18°C and *X.tropicalis* at 25°C with a light/dark cycle of 12 h/12 h.

For this study, we selected a total of n=96 samples, representing the development of *Xenopus*, from neurula to mid-life adult stages: 11 samples from peripheral blood, 11 samples from brain, 11 samples from liver, 11 samples from hind limb thigh muscle, 11 samples from skin, 17 samples from toes, and 24 samples from whole embryos and juveniles NF stage 18, 28, 47, 55, 58 and 66 *X. laevis* and stage 18 and 28 *X. tropicalis*. 31 outlier samples from a variety of tissues from both *Xenopus* species were excluded from analysis on the basis of the DNAm profile. Animals were anesthetized in 0.15% MS-222 (Sigma-Adrich, A5040) and sacrificed by transection between the brainstem and the spinal cord. After harvesting, the tissues were snap frozen in liquid nitrogen and grinded to powder using a mortar and a pestle. For genomic DNA extraction, 25 mg tissue were mixed with 20 μl 20 mg/ml Proteinase K (Qiagen, 19131) and 180 μl buffer ATL (Qiagen, 19076) and incubated for 1 h at 56°C, 1000 rpm at an Eppendorf Thermomixer Comfort. For genomic DNA extraction from blood, we used 10 μl peripheral blood. DNA extraction was performed using the DNeasy Blood & Tissue kit (Qiagen, 69504). DNA was eluted in 100 μl buffer AE (10 mM Tris-Cl, 0.5 mM EDTA; pH 9.0). Samples with DNA concentration below 50 ng/μl were concentrated by ethanol precipitation. The DNA pellet was washed with 70% ethanol and resuspended in 50 μl AE buffer.

### Human tissue samples

To build the human-clawed frog clock, we analyzed previously generated methylation data from n=1366 human tissue samples (adipose, blood, bone marrow, cerebellum, cortex, dermis, epidermis, embryonic cells, fibroblasts, heart, keratinocytes, kidney, liver, lung, lymph node, muscle, pituitary, placenta, skin, spleen) from individuals whose ages ranged from 0 to 101 years [51]. The tissue samples came from three sources. Tissue and organ samples were from the National NeuroAIDS Tissue Consortium [52]. Blood samples were from the Cape Town Adolescent Antiretroviral Cohort study [53]. Skin and other primary cells were provided by Kenneth Raj [54]. Ethics approval (IRB#15-001454, IRB#16-000471, IRB#18-000315, IRB#16-002028).

### DNA methylation data

All DNA methylation data were generated using the custom Infinium array ‘HorvathMammalMethylChip40” [15]. By design, the mammalian methylation array facilitates epigenetic studies across mammalian species (including humans) due to its very high coverage (over thousand-fold) of highly-conserved CpGs in mammals. A subset of cytosines on the mammalian array also apply to more distant species including amphibians [15]. Only 4,635 CpGs out of all CpGs on the mammalian array actually map to one or both of the clawed frog genomes (1,829 map to African clawed frog, 4,239 map to Western clawed frog) according to genome assemblies XenTro9.1.102 and XenLae10.1 from ENSEMBL. Genome coordinate information can be downloaded from our GitHub page (https://github.com/shorvath/MammalianMethylationConsortium) and the supplementary information in [15]. The chip manifest file can be found at Gene Expression Omnibus (GEO) at NCBI as platform GPL28271. The SeSaMe normalization method was used to define beta values for each probe [55].

### Penalized Regression models

Details on the clocks (CpGs, genome coordinates) and R software code are provided in **Suppl. Table 1**. Penalized regression models were created with glmnet [56]. We investigated models produced by “elastic net” regression (alpha=0.5). The optimal penalty parameters in all cases were determined automatically by using a 10-fold internal cross-validation (cv.glmnet) on the training set. By definition, the alpha value for the elastic net regression was set to 0.5 (midpoint between Ridge and Lasso type regression) and was not optimized for model performance.

We performed a cross-validation scheme for arriving at unbiased (or at least less biased) estimates of the accuracy of the different DNAm based age estimators. For validation of the clocks, we used leave-one-out LOO cross-validation (LOOCV) in which one sample was left out of the regression, then predicted the age for the remaining samples and iterated this process over all samples.

A critical step is the transformation of chronological age (the dependent variable). While no transformation was used for the single-species pan-tissue clocks for *X.laevis* and *X.tropicalis*, respectively, we did use a log linear transformation for the 2-species pan-tissue clocks for clawed frogs and for the dual species clock of chronological age (Supplement).

### Relative age estimation

To introduce biological meaning into age estimates of two clawed frog species and humans that all have very different lifespan; as well as to overcome the inevitable skewing due to unequal distribution of data points from clawed frogs and humans across age range, relative age estimation was made using the formula: Relative age = Age/maxLifespan, where the maximum lifespan for the three species was chosen from the *anAge* database [44]. Maximum age of African clawed frogs and Western clawed frogs was 30.3 and 16 years, respectively, and the maximum age of humans was 122.5 years. The oldest frog *X.tropicalis* (16 years) is currently (July 2022) still alive in the lab of Christof Niehrs.

### Epigenome wide association studies of age

EWAS was performed in each tissue and frog species separately with the R function “standardScreeningNumericTrait” in the “WGCNA” R package [57]. Next, the results were combined across tissues with Stouffer’s meta-analysis method and combined across species.

### GREAT functional enrichment analysis

We used the Genomic Regions Enrichment of Annotations Tool (GREAT) to analyze the age related CpGs [58]. This software tool has not yet been adapted for frogs. Rather, we used the human hg19 genome assembly. To avoid biases arising from the use of the mammalian array platform, we restricted the background according to the genomic regions covered by the 4,635 probes that mapped to the *X.tropicalis* genome. GREAT calculates statistics by associating genomic regions with nearby genes and applying the gene annotations to the regions. Association is a two-step process. First, every gene is assigned a regulatory domain. Then, each genomic region is associated with all genes whose regulatory domain it overlaps. To define the gene regulatory domain, each gene is assigned a basal regulatory domain of a minimum distance upstream and downstream of the TSS (regardless of other nearby genes). We used the settings: Proximal: 5.0 kb upstream, 1.0 kb downstream, plus Distal: up to 50 kb). Gene set enrichment was done for gene ontology, molecular pathways, diseases, upstream regulators, human and mouse phenotypes.

### Genome annotation

We aligned microarray probes to the reference genomes of Xenopus_tropicalis 9.1.102 and Xenopus_laevis 10.1 from ENSEMBL. The alignment was done using the QUASR package [59], with the assumption of bisulfite conversion treatment of the genomic DNA. Following the alignment, the CpGs were annotated based on the distance to the closest transcriptional start site using the ChIPseeker package [60] (**Suppl. Table 2**).

## Results

### Two frog species

We generated novel DNA methylation data and characterized DNAm from *X. laevis* (n=59 samples) and *X. tropicalis* (n=37 samples). We profiled six tissues (blood, brain, skin, lung, muscle, toe) as well as whole embryos, tadpoles, and juveniles, spanning a wide age range from 2-day-to 19-year-old whole animals and tissues, respectively (**Tables 1-3**). Since we were interested in comparing frogs to humans and other mammals, we used the mammalian methylation array platform that profiles individual CpGs in highly conserved stretches of DNA in mammals [15]. By design, the mammalian methylation array facilitates epigenetic clock studies across mammalian species (including humans) due to its very high sequencing depth (over thousand-fold) in highly-conserved CpGs in mammals. Only 4,635 CpGs out of 37492 CpGs on the mammalian array map to one or both clawed frog genomes (1,829 CpGs map to *X.laevis*, 4,239 CpGs map to *X.tropicalis*) according to genome assemblies XenTro9.1.102 and XenLae10.1 from ENSEMBL.

**Table 1.** Description of *X. laevis* data by tissue type.

| Tissue | N | Mean Age | Min. Age | Max. Age |
| --- | --- | --- | --- | --- |
| Blood | 6 | 9.42 years | 9 months | 19 years |
| Brain | 6 | 9.42 years | 9 months | 19 years |
| Lung | 6 | 9.42 years | 9 months | 19 years |
| Muscle | 6 | 9.42 years | 9 months | 19 years |
| Skin | 6 | 9.42 years | 9 months | 19 years |
| Toe | 9 | 7.83 years | 9 months | 19 years |
| Whole Animal | 20 | 3.2 months | 2 days | 7 months |

N=Total number of samples. Age: mean, minimum and maximum.

**Table 2.** Description of *X. tropicalis* data by tissue type.

| Tissue | N | Mean Age | Min. Age | Max. Age |
| --- | --- | --- | --- | --- |
| Blood | 5 | 6.2 years | 2 years | 10 years |
| Brain | 5 | 6.2 years | 2 years | 10 years |
| Lung | 5 | 6.2 years | 2 years | 10 years |
| Muscle | 5 | 6.2 years | 2 years | 10 years |
| Skin | 5 | 6.2 years | 2 years | 10 years |
| Toe | 8 | 6 | 2 years | 10 years |
| Whole Animal | 4 | 2.5 days | 2 days | 3 days |

N=Total number of samples. Age: mean, minimum and maximum.

**Table 3.**

| Tissue | N | Dev. stage | Age |
| --- | --- | --- | --- |
| *X. laevis* | 2 | 18 | 2 days |
|  | 3 | 28 | 3 days |
|  | 5 | 47 | 9 days |
|  | 8 | 55 | 7 months |
|  | 12 | N/A | 9 months |
|  | 1 | N/A | 2 years |
| *X. tropicalis* | 2 | 18 | 2 days |
|  | 2 | 28 | 3 days |
|  | 13 | N/A | 2 years |

N=Total number of samples. Age: Developmental stage, chronological age.

Unsupervised hierarchical clustering of the frog tissue samples led to two insights (**Fig. 1**). First, a large number of samples are putative outliers (n=31 out of 96) that do not fall into specific clusters/branches at a lenient height cut-off of 0.25 (Figure 1). To err on the side of caution, we removed these 31 putative outliers from the rest of the analysis. Second, the non-outlying samples fall into clusters that correspond to species and to a lesser extent to tissue type.

**Figure 1.**
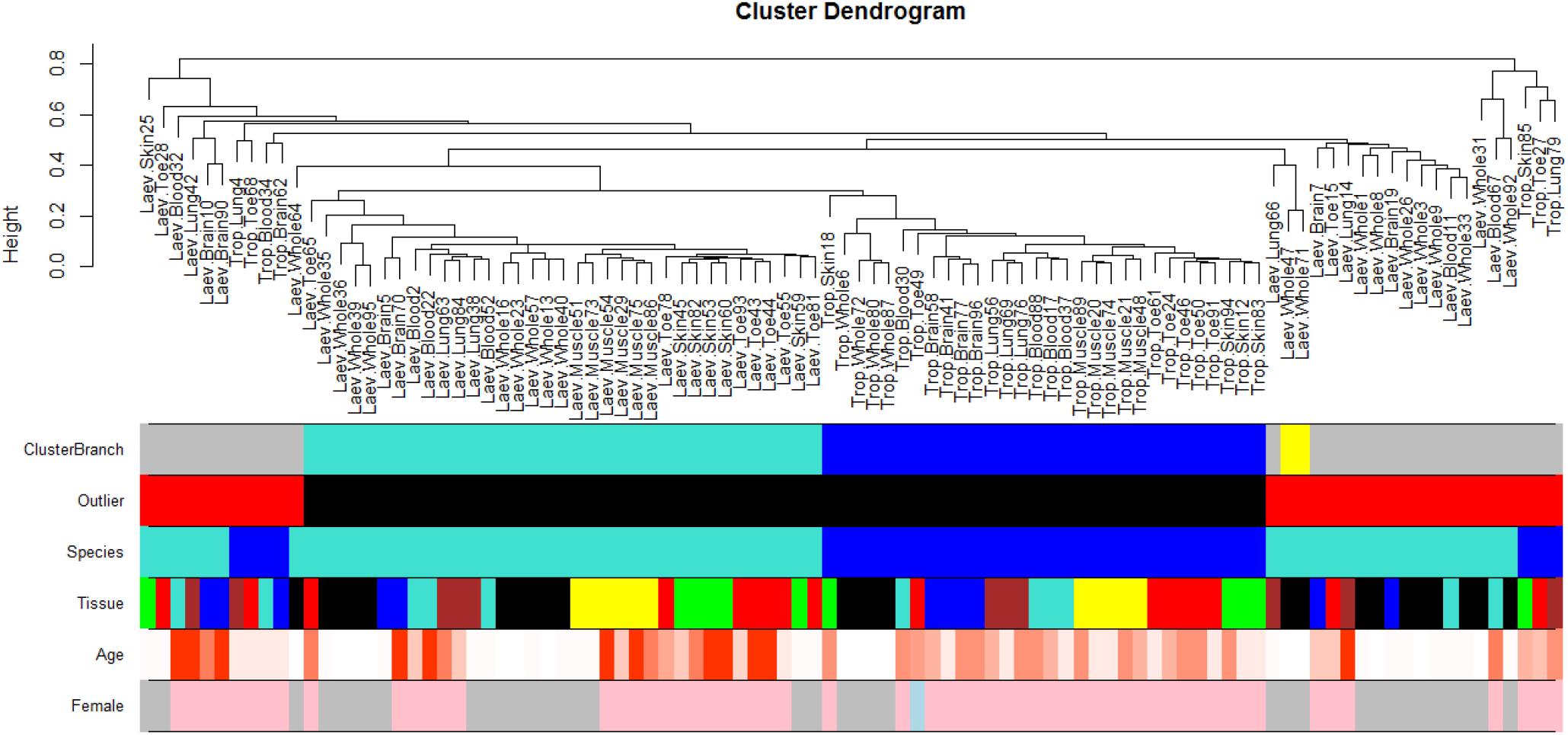
Unsupervised hierarchical clustering of *Xenopus* tissues. The clustering height (y-axis) can be interpreted as distance based on pairwise correlation coefficients. Color bands underneath indicate clustering branch (corresponding to a height cut-off of 0.29 on the y-axis), outlier status (red=outlier), frog species (blue=tropicalis, turquoise=laevis), tissue type (see the labels), age (red corresponds to old age), and sex (pink=female, blue=male, grey=unknown). As dissimilarity, we used 1 minus the Pearson correlation coefficient across the 4635 CpGs that map to tropicalis and/or laevis. We used average linkage as an intergroup dissimilarity measure.

### Epigenetic clocks

We used these high quality DNAm data to construct different epigenetic clocks for clawed frogs only and for both human and clawed frogs. For the construction of the dual species human-clawed frog clock, we used the DNAm data previously generated with the HorvathMammalMethylChip40 in 1366 human samples representing 20 tissues from individuals 0 to 101 years old [51]. Our *Xenopus* clocks can be distinguished along three dimensions, species, age range, and measure of age. We used a combined set of all samples to train a pan-tissue clock (pan-clock) suited for age predictions across different tissue types included in the clock construction. We also created clocks tailor-made for specific species, which were trained based on all samples from the *laevis*-clock and the *tropicalis*-clock. We also created a pan-tissue clock for young organisms (young-lock) trained on samples coming from animals no older than two years of age.

While the pan-tissue *Xenopus* clocks apply only to clawed frogs, we also created dual-species clocks, referred to as human-clawed frog clocks, for estimates of chronological age and relative age. Relative age is the ratio of chronological age to maximum lifespan (i.e., the maximum age of death observed in the species). Thus, relative age takes on values between 0 and 1. The maximum lifespan observed for *X.laevis* and *X.tropicalis* was 30.3 and 16 years, respectively, and the maximum lifespan observed for humans was 122.5 years. Relative age allows for alignment and biologically meaningful comparison between species with different lifespan (clawed frogs and humans), which is not afforded by mere measurement of chronological age.

To arrive at unbiased estimates of the epigenetic clocks, we used leave-one-out (LOO) cross-validation of the training data. The cross-validation study reports unbiased estimates of the age correlation R (defined as Pearson correlation between the age estimate (DNAm age) and chronological age) as well as the median absolute error (MAE) measuring the deviation between the predicted and observed age (for chronological age in years). As indicated by their names, the pan-clock is highly accurate in age estimation of the different tissue samples (R=0.88 and median error 2.05 years, **Fig. 2A**), and the young-clock is highly accurate in age estimation of the different tissue samples coming from young animals (R=0.93 and median error 0.107 years, **Fig. 2D**). We also developed highly accurate clawed frog clocks for single species: *X.laevis* (R=0.91 and median error 3.24 years, **Fig. 2B**) and *X.tropicalis* (R=0.96 and median error 0.946 years, **Figure 2C**).

**Figure 2.**
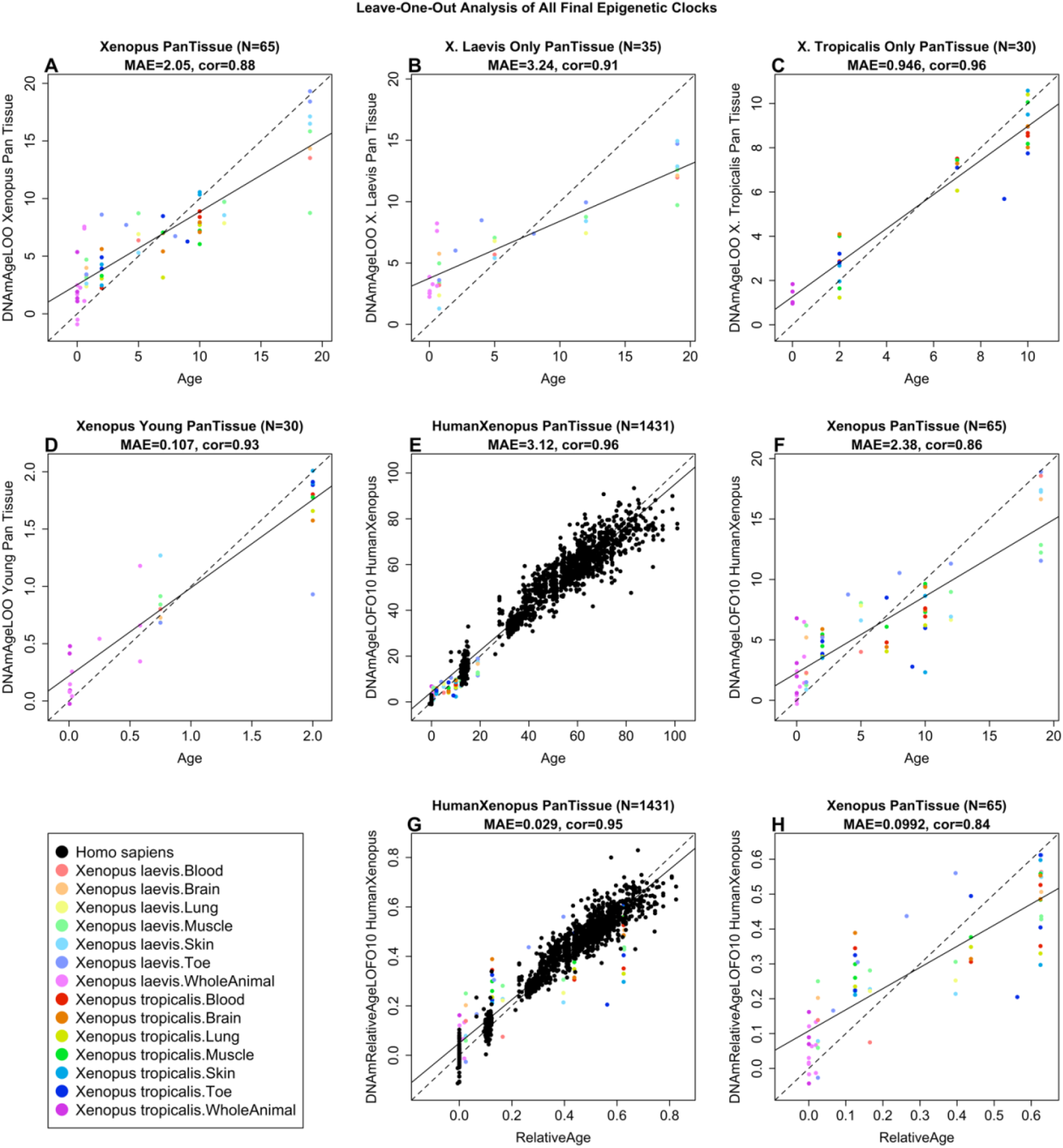
Cross-validation study of epigenetic clocks for *Xenopus* and humans. (**A-D**) Four epigenetic clocks that only apply to *Xenopus*. Leave-one-sample-out estimate of DNA methylation age (y-axis, in units of years) versus chronological age for **A**) all *Xenopus* tissues (both species), **B**) all tissues from *X.laevis*, **C)** all tissues from *X. tropicalis*, **D**) all tissues from young *Xenopus* (age <= 2 years). Ten-fold cross validation analysis of the human-clawed frog clocks for **E-F**) chronological age and **G-H**) relative age, respectively. **E,G**) Human samples are colored in black and *Xenopus* samples are colored by species and tissue type, and analogous in **F,H**) but restricted to *Xenopus* samples (colored by *Xenopus* tissue type). Each panel reports the sample size (in parenthesis), correlation coefficient, median absolute error (MAE).

We developed two dual-species clocks based on our clawed frog samples and previously characterized human tissues. The interest to create such dual-species clocks is (i) that they are expected to increase the likelihood that findings in one species will translate to the other and (ii) to increase statistical significance. The human-clawed clock for chronological age (R=0.96 for the human and clawed frog samples and R=0.86 for the clawed frog samples, **Fig. 2E-F**) and relative age (R=0.95 for the human and clawed frog samples and R=0.84 for the clawed frog samples, **Fig. 2G-H**).

### Epigenome-wide association study of age

We performed three separate epigenome-wide association studies (EWAS) of age which involved (i) *X.laevis* only (n=35), (ii) *X.tropicalis* only (n=30), and iii) Stouffer meta-analysis combining the former two EWAS results (**Fig. 3A-C; Suppl. Table 2**). Due to the low sample size, we ignored tissue type in this analysis. Each EWAS correlated chronological age with individual CpG methylation levels. The analysis was restricted to 4,239 CpGs that mapped to the genome of *X.tropicalis* (XenTro9.1.102).

**Figure 3.**
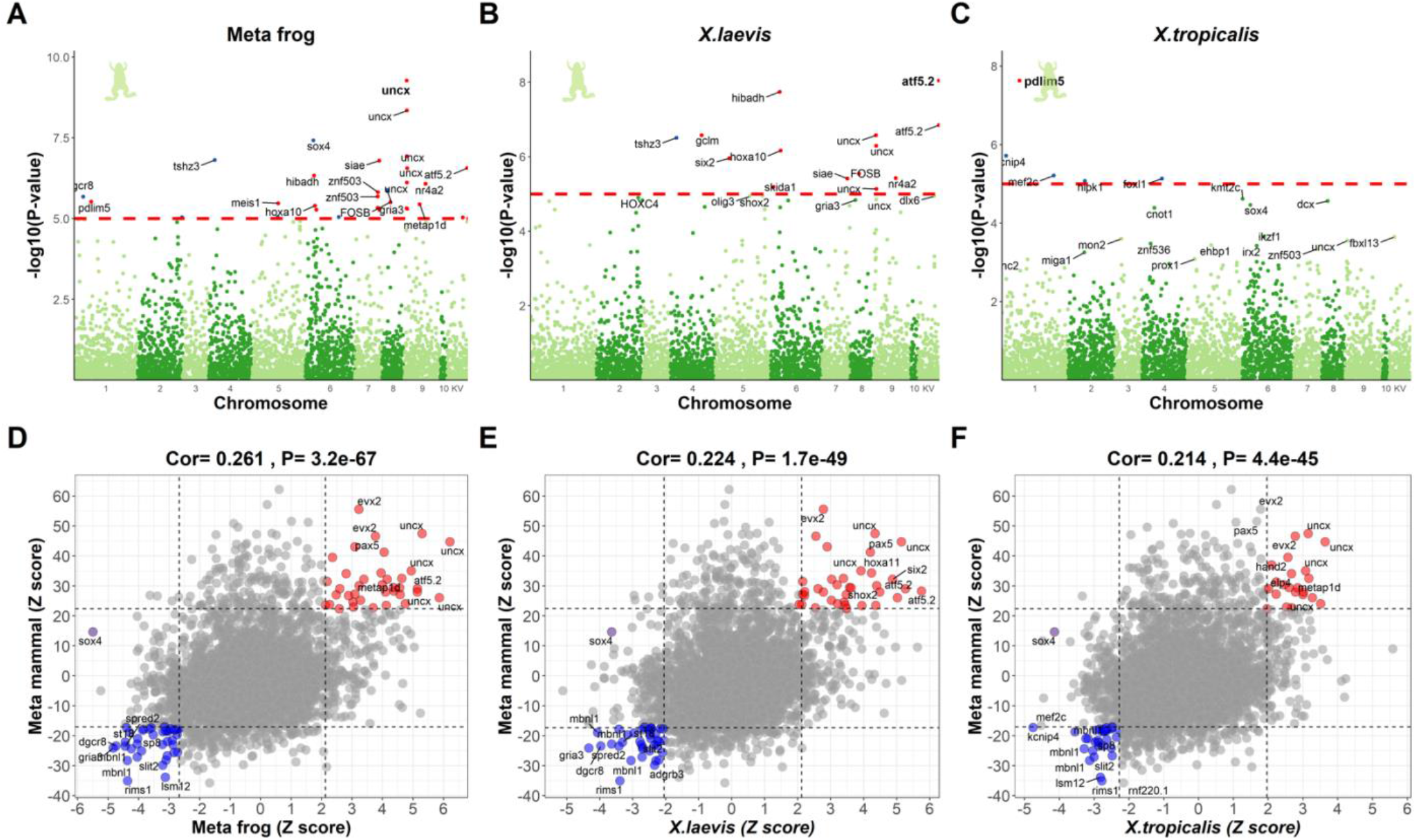
EWAS of age in *Xenopus*. The top panels display Manhattan plots for EWAS of age (**A**) Stouffer’s method meta-analysis that combines (**B**)*X.laevis*, and (**C**)*X.tropicalis* across all tissue types. The red dash lines indicate suggestive levels of significance at P<1.0E-05. CpGs are colored in red or blue for positive or negative age correlations, respectively. The y-axis displays -log base 10 transformed P value and the x-axis displays chromosome number based on the *X. tropicalis* genome (v9.1.102). Chromosome KV denotes the alias names for the CpGs with unspecified chromosomes. (**D-F**) Display the scatter plots between frog EWAS of age and Eutherian EWAS of age, based on Z statistics. The title lists the number of CpGs (4,239) in our mammalian array that can map to the *X. tropicalis* genome and the Pearson correlation and its P-value between the two EWAS Z statistics. Each dot corresponds to a CpG. The CpG cg17865363 exhibiting highly significant P value in frog EWAS is annotated by its nearby gene *sox4* and marked in purple. Labels are provided for the top 10 hypermethylated/hypomethylated CpGs according to the product of Z scores in x and y-axis.

At a genome wide significance level of P<1.0×10-7, two CpGs were significant in the *laevis* EWAS (**Fig. 3B; Suppl. Table 2**). One CpG cg10071691 (P=9.1×10^-9^) is located near *atf5.2* on an unspecified chromosome (alias name: KV460357.1). The other one, cg11266179 (P=1.8×10^-8^), is located near *hibadh*. In tropicalis, only one CpG (P=2.3×10^-8^) near *pdlim5* reached genome wide significance (**Fig. 3C; Suppl. Table 2**).

To combine the EWAS results of both *Xenopus* species, we used Stouffer meta-analysis method (equal weights) resulting in a Z statistic that follows a standard normal distribution under the null hypothesis of zero correlation with age. Comparing the frog EWAS results with those from the Mammalian Methylation Consortium reveals that CpGs in the *UNCX* gene are also highly correlated with chronological age in many mammalian tissues as shown in our pan-mammalian ageing studies [16]. The EWAS of age results in frogs correlated positively (Pearson correlation r=0.26) with the EWAS of age results in eutherian mammalian species (**Fig. 3D; Suppl. Table 2)** revealing an age-related gain of methylation in genes that play a role notably in neural development (e.g., *uncx, sox4, pax5* and *evx2*). Similar correlation with the EWAS results from mammals could be observed when focusing the analysis on EWAS findings from a single frog species (*r* > 0.2**, Fig. 3E-F**).

*Uncx* was the top gene with positive CpG methylation-age correlation. Of the 4239 CpGs, 10 significantly age-related CpGs located near *uncx*, mostly in the promoter (**Fig. 3A, Suppl. Table 2**). Human *UNCX* encodes a homeobox transcription factor whose mouse homolog is implicated the development of the axial skeleton and in neural progenitor regulation in the olfactory epithelium [61, 62] and for which increased DNAme (cg04816311) is associated with target organ damage in older African Americans [63]. Notably, mutations in the *uncx* orthologue in C. elegans, *unc4*, extends male lifespan [64]. A positive age-related CpG was found near *hibadh*, where CpG methylation gain of human *HIBADH* (cg01065605) positively correlates with mortality in the InCHIANTI cohort [65]. For *nr4a2*, downregulation of the respective rat gene was reported in the hippocampus of aged animals [66].

The top ranking negative age-related CpG is located near *sox4* (*SRY-box transcription factor 4*), which regulates retinal precursor development in *Xenopus* [67]. DNAme or expression of *SOX4* in mice and humans are associated with ageing or age-related disease, notably cancer [68, 69]. A negative age-related CpG is located near *tshz3 (Teashirt Zinc Finger Homeobox 3*), rare protein-altering variants of which are highly enriched in a cohort of supercentenarians [70]. It is noteworthy that negative age-related CpG associated gene orthologs tend to be involved in synaptic transmission, including *tshz3, dgcr8, spred2, gria3, gria4, kcnip4*, and *rims1* [71–76]. Indeed, the term ‘synaptic transmission’ was retrieved in GREAT functional enrichment analysis among the genes that lose methylation with age (**Fig. 4**). Once again, these genes are also associated with negative age-related CpGs in mammals [16, 77, 78].

**Figure 4.**
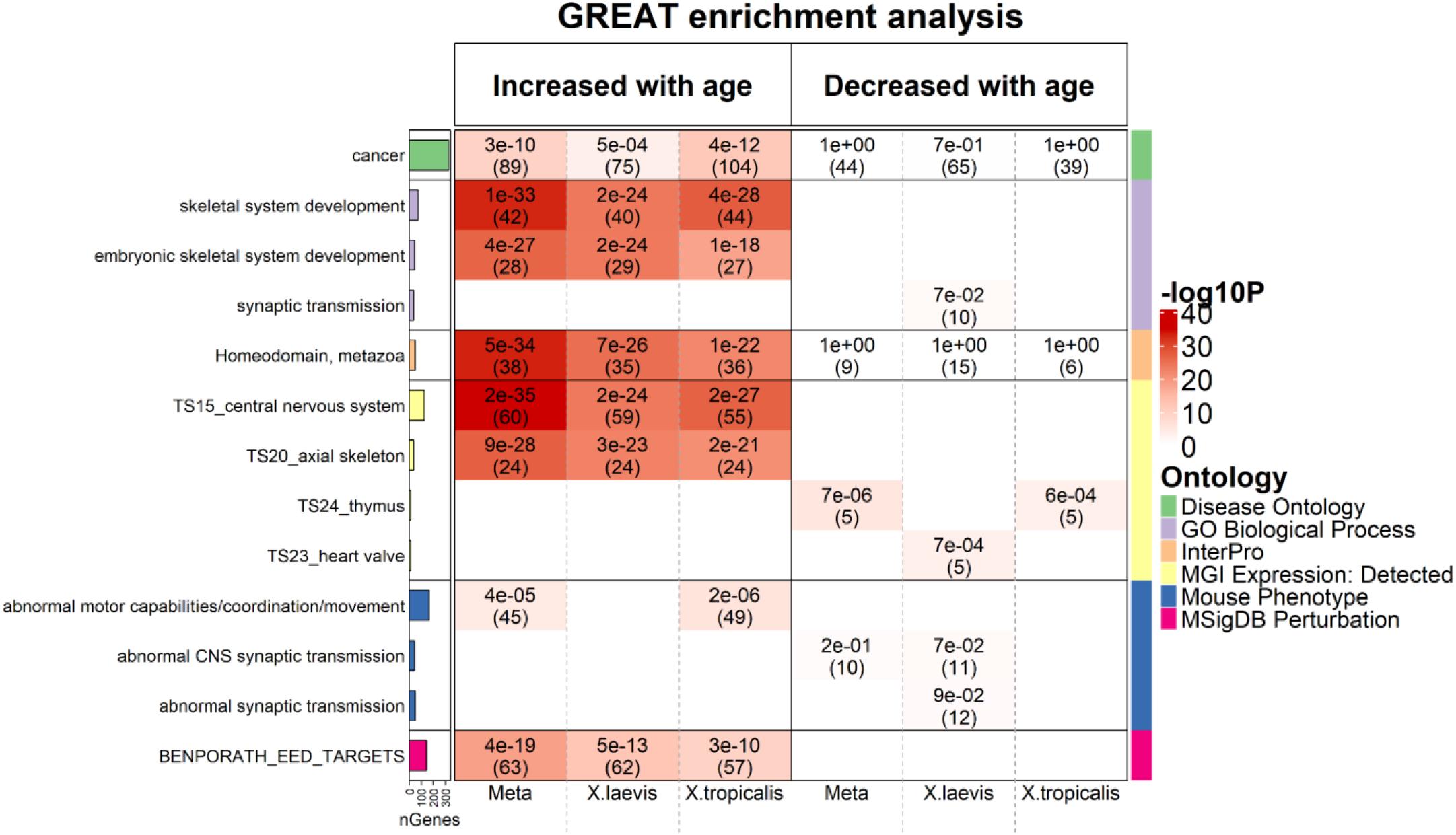
Genomic region-based GREAT functional enrichment analysis. GREAT functional enrichment analysis was based on the top 500 CpGs that increased or decreased with age from EWAS in (1) meta-analysis, (2) *X.laevis*, and (3)*X.tropicalis*, respectively (**Suppl. Table 2**). The background was based on the genomic regions of the 4,239 mammalian CpGs and the assembly in hg19. The y-axis lists the name of a functional gene set/biological pathway, sorted by ontology and the most significant hypergeometric P value within each ontology. The bar plots in the first column report the total number of genes at each studied gene set adjusted based on our background. The left and right panels of the x-axis list the enrichment results based on the top 500 CpGs with positive and negative age correlation. We list unadjusted hypergeometric P value (number of overlap genes) at each cell, provided P<0.1. The heatmap color-codes -log10 (P-value). Abbreviations: BENPORATH_EED_TARGETS denotes “EED targets: genes identified by ChIP on chip as targets of the Polycomb protein EED (GeneID=8726) in human embryonic stem cells.”

### Functional enrichment of EWAS of age

To annotate the biological functions of the age-related CpGs in frogs, we performed genomic region GREAT functional enrichment analysis [58]. The functional annotations were based on the top 500 positive and the top 500 negative age-related CpGs from each of the 3 EWAS studies. The background underlying the GREAT analysis was based on the genomic regions of the 4239 CpGs in Hg19 assembly. Choosing this background set of CpGs ensured that our enrichment analyses were not biased by the content/design of the mammalian array.

The positive age-related CpGs were enriched in several gene sets involved in developmental processes, notably embryonic skeletal system development in GO term (GREAT P-value <1.0×10^-33^), target sites of Polycomb repressive complex 2 (PRC2) such the subunit EED in the MSigDB database (P-value <1.0×10^-19^, **Fig. 4**). Similarly, we find enrichments for gene sets that play a role in the development of mice including axial skeleton, and skeleton or rib morphology (**Fig. 4**). Moreover, ‘homeodomain’ and genes expressed in mouse central nervous system were highly enriched terms (GREAT P-value 5×10^-34^ and 2×10^-35^, respectively).

Fewer significant enrichments can be observed for negatively age-related CpGs. The identified gene sets include GO terms such as regulation of synaptic transmission, RNA splicing, and genes expressed in the thymus or heart valve according to MGI expression, regulation of RNA splicing under GO term (**Fig. 4 and Supplementary Figure S1**).

### Human chromatin state annotation

From the outset, it makes little sense to use a *human* chromatin state analysis to interpret the EWAS of age results in frogs. However, we present these results since the findings turn out to be consistent with those from the EWAS of age by the Mammalian Methylation Consortium [16] as detailed in the following. We used universal human chromatin states based on 1,032 experiments that mapped 32 types of chromatin marks in over 100 human cell and tissue types [79]. First, we used the hypergeometric test-based overlap analysis between chromatin states and the top 500 *positively* age related CpGs. Again, the analysis properly adjusted for the background set of CpGs that map to the frog genome and the array platform.

For the positively age related CpGs, we observed significant overlap with bivalent regulatory regions (specifically bivalent promoter 2 and bivalent promoter 4) and a Polycomb repressed state (ReprPC1). These three states contain a high proportion of CpGs at target sites of Polycomb repressive complex 2 (PRC2). The frog meta-analysis EWAS of age exhibits significant overlap with BivProm2 state with an odds ratio greater than 2 (hypergeometric P=9.1×10^-4^, **Suppl. Fig. S1**). In contrast, the negatively age-related CpGs (top 500) are enriched in select weak enhancer state EnhWk4, enhancer state EnhA2 and transcribed exon state TxEx4.

### Overlap with PRC2 target sites

Since a hallmark of positive age-related CpGs in mammals is their association with regions that are targeted by PRC2 [2, 80, 81], we further examined the overlap between the age-related CpGs and target sites of both PRC1 and PRC2. Toward this end, we used PRC binding annotations from human cells. We defined PRC annotations based on the binding of at least two transcription factor members of PRC1 (subunits: RING1, RNF2, BMI1) or of PRC2 (subunits: EED,SUZ12 and EZH2) in 49 ChipSeq datasets available in ENCODE [82]. The top 500 CpGs with positive age correlation are enriched in PRC2 target sits based on the meta EWAS (hypergeometric P-value =7.8×10^-35^ and odds ratio (OR)=4.4), *X.laevis* EWAS (OR=3.3 and P=5.5×10^-22^) and *X.tropicalis* EWAS (OR=3.9 and P=3.0×10^-29^) (**Suppl. Fig. S1**). By contrast, the overlap with PRC1 target sites is far less significant (**Suppl. Fig. S1**). The results suggest that the association of ageing-related methylated CpGs with PRC2 target genes is evolutionary conserved between mammals and amphibians.

## Discussion

The key finding of this study is the evolutionary conservation of epigenetic ageing signatures between frogs and humans. This conservation relates to (i) our ability to construct two epigenetic clocks that apply to *Xenopus* and humans, (ii) that age-related CpGs are located near genes also found in mammalian clocks and that are implicated in age-associated disease, (iii) that positive age-related CpGs are associated with PRC2 target sides, a hallmark also observed in mammals.

Our study has several limitations. First, 31 out of 96 tissue samples were deemed to be putative outliers according to our unsupervised hierarchical clustering analysis. The large number of outliers unlikely reflects DNA quality issues since DNA quality and quantity the DNA was carefully assessed by Qubit analysis of sufficient quality and quantity according to standard laboratory assessments (Qubit). Instead, we believe that the high number of outlying data reflects the fact that the mammalian array is either imperfectly suited for DNA samples from amphibians or that the aliquoting of frozen-ground whole juveniles introduced inhomogeneities in tissue representation. Second, the mammalian array profiles only about 4239 CpGs out of millions of CpGs in the tropicalis genome. While the large sequencing depth at highly conserved CpGs is ideal for building human-frog clocks, the low number of cytosines will be limiting for studies that aim to characterize regulatory relationships between methylation and transcriptomic changes.

A striking finding of this study is the construction of two epigenetic clocks that apply to *Xenopus* and humans. The two clocks have different interpretations: the first clock measures chronological age in both species. The second clock measures relative age (age divided by maximum lifespan). Relative age may be a biologically more meaningful measure since it adjusts for the strong difference in lifespan. Each of these dual-species clocks estimates age based on a single mathematical formula derived from a multivariate regression model focusing on highly conserved CpGs. The fact that we could successfully construct human-clawed frog clocks is due to both biological and technical reasons. A biological reason is the high conservation of (positively) age related changes in PRC2 target sites, as can be seen from our EWAS of age. A technical reason is the large sequencing depth at highly conserved CpGs that were profiled on the mammalian methylation array platform [15]. Our dual species human-clawed frog clocks, for absolute and relative age, increase the chance that findings in frogs translate to humans and vice versa. The bias due to differences in maximum lifespan is mitigated by the generation of the human-clawed frog clocks for *relative* age, which embeds the estimated age in context of the maximal lifespan recorded for the relevant species. The high accuracy of these clocks demonstrates that one can build epigenetic clocks for two species based on a single mathematical formula. Treatments that alter the epigenetic age in *Xenopus* are therefore likely to exert similar effects in humans.

We also present *Xenopus* clocks that were trained in *laevis* and *X. tropicalis*. We generated DNA methylation data from six tissue types and from whole animals in these two species. Using these DNAm data, we trained and validated highly accurate age estimators (epigenetic clocks) that apply to the developmental life course (from birth to mid-life), and identified genes associated with the ageing process in the clawed frog. These data allowed us to construct a highly accurate pan-tissue age estimator (pan-clock) based on six clawed frog tissue types (blood, brain, liver, muscle, skin, toe) and whole animal, and clocks developed based on individual *Xenopus* species, as well as a clock developed based on all tissues from embryos and juveniles (young-clock). Given that all four pure *Xenopus* clocks can estimate age in six different tissues, we anticipate that they apply to additional tissues as well. However, we cannot rule out that these clocks could fail in some highly specialized cell types. We expect that the *Xenopus* clocks apply to other clawed frog species as well in the sense that they will lead to high age correlations in tissue samples collected from both young and old frogs. However, epigenetic age predictions can differ from the true chronological age by a constant offset/bias in new tissue types or new frog species due to biological and technical reasons including differences in probe sequence conservation or storage conditions. The offset/bias can be estimated from the data by including tissue samples from frogs of known ages.

These epigenetic clocks reveal several salient features with regard to the biology of ageing. First, the *Xenopus* pan-tissue clock reaffirms the conclusion drawn from the human pan-tissue clock, which is that ageing might be a systemic biological process that affects the whole body. Second, the ability to combine these two pan-tissue clocks into a single human-clawed frog pan-tissue clock, species whose lineages diverged some 350 million years ago (according to timetree.org) [83], attests to the high conservation of the ageing process. This conclusion is corroborated by our EWAS analysis. Previous studies in humans showed that a hallmark of age-related CpGs is their association with target sites of PRC2, which gain methylation with age [2, 80, 81] and this feature is fully recapitulated in *Xenopus*. The physiological significance of this association is an important open question. PRC2 plays a prominent role during embryonic development [84] and consequently many ageing-clock-associated genes relate to developmental processes. Given its evolutionary conservation from frogs to human, the methylation status of PRC2 targets supports some critical causal relationship to systemic ageing. Since the association with PRC2 with ageing stems from analyses of adult, postmitotic cells and of different tissue origin rather than from embryonic cells, is tempting to speculate that the adult methylation status will get important input during embryonic development, the very phase when PRC2 target gene expression is prominent. Indeed, according to *Xenopus* epigenetic clocks, epigenetic ageing proceeds already during embryonic and larval development, long before metamorphosis, which only begins weeks after fertilization. Consistent with the idea of ‘embryonic ageing’, *Xenopus* genes associated with positive age-related CpGs encode many developmental regulators. In particular, it is noteworthy that genes associated with both positive and negative age-related CpGs relate to neural processes, although in somewhat opposite direction: While DNAme increase is linked to neural developmental genes, DNAme decrease links to synaptic transmission, roughly corresponding to processes of immature- *vs*. mature neuronal cells, respectively. Altogether, this leads to the counter-intuitive suggestion that studying *Xenopus* neural development may yield new insights into biological ageing. The availability of epigenetic clocks as quantitative, accurate age biomarker overcomes the limitations set by the decade lifetimes of clawed frogs and render this endeavor feasible.

## Supporting information

Supplemental Table 1. Frog Clocks

Supplementary Methods. Software for Xenopus clocks.

Supplementary Table 2. EWAS of age in Xenopus. Human Genome Coordinates HG38

## Acknowledgements

This work was supported by the Paul G. Allen Frontiers Group (PI SH) and Open Philanthropy (SH). We thank the European *Xenopus* Resource Centre (EXRC) for DNA samples.

## Data availability

The data will be made publicly available on Gene Expression Omnibus as part of the data release from the Mammalian Methylation Consortium. Genome annotations of these CpGs can be found on GitHub https://github.com/shorvath/MammalianMethylationConsortium. The mammalian methylation array is broadly available to the research community from the non-profit Epigenetic Clock Development Foundation (https://clockfoundation.org/).

## Author contributions

CN and EP provided *Xenopus* tissue samples. JAZ, ATL, AH, and SH carried out the statistical analysis. JAZ, ATL, EP, SH, CN wrote the first draft of the article. CN and SH conceived and supervised the study.

---

**Supplementary Figure S1.**
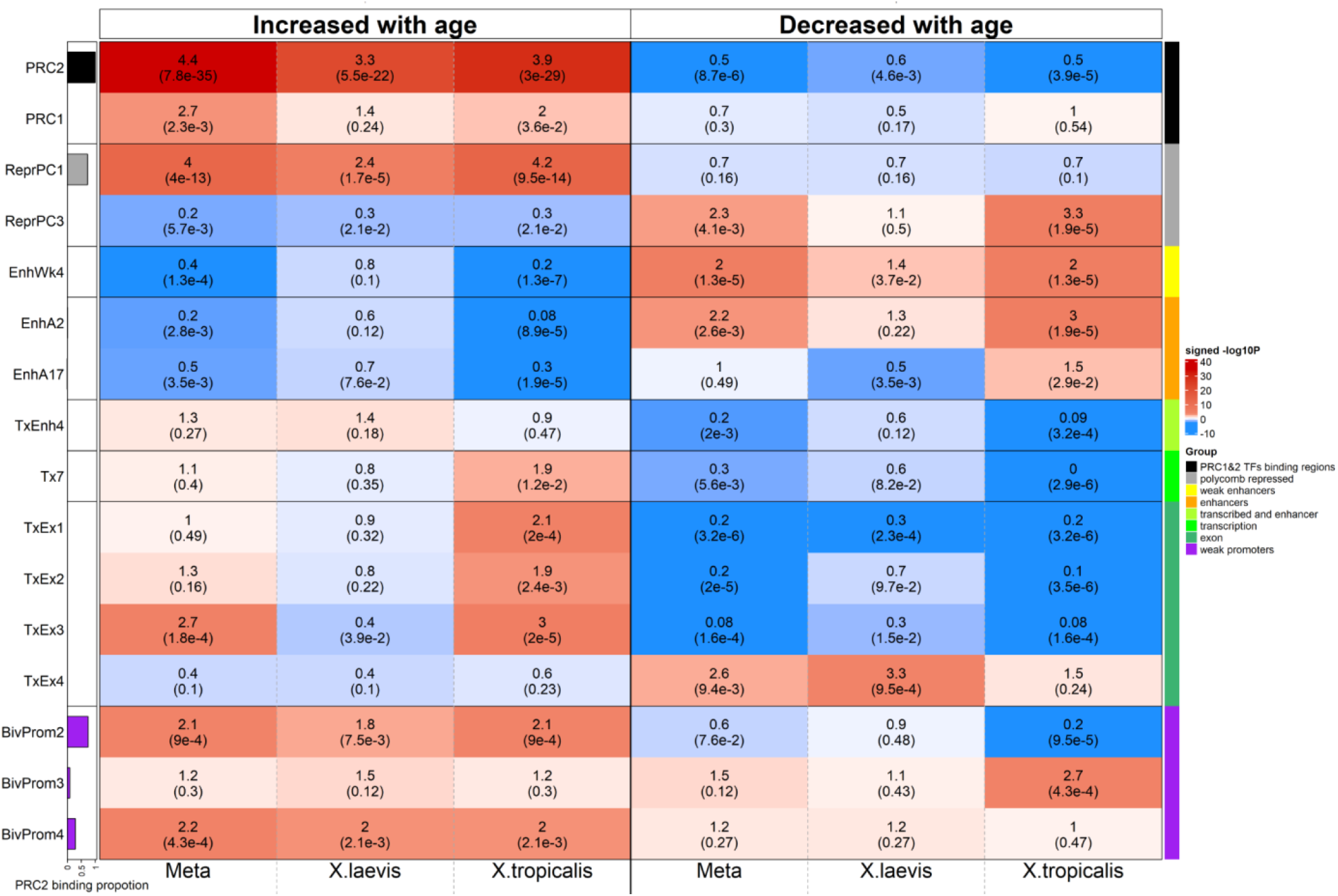
Chromatin state analysis of age-related CpGs. The heatmap color-codes the hypergeometric overlap analysis between age-related CpGs (columns) and two groupings of CpGs (a) binding by Polycomb repressive complex 1 and 2 (PRC1, PRC2) defined based on ChipSeq datasets in ENCODE [82] and (b) universal chromatin states analysis [79], see the first two rows. The background is based on the 4,239 mammalian CpGs that can map to the *X.tropicalis* genome (v9.1.102). The first column shows a bar plot that reports the proportion of CpGs bound by PRC2 that ranges from zero (RPC1) to one (PRC2). For each row (chromatin state or PRC annotation), the table reports odds ratios (OR) from hypergeometric test results for the top 500 CpGs that increased/decreased with age from meta-EWAS, *X.laevis* EWAS and *X.tropicalis* EWAS, respectively. The heatmap color gradient is based on -log10 (unadjusted hypergeometric P value) multiplied by the sign of OR greater than one. Red colors denote OR greater than one in contrast with blue colors for OR less than one. Legend lists states based on their group category and PRC group. The y-axis lists chromatin states and PRC2 target sites. The left/right panel lists the results based on the top 500 CpGs with positive/negative age correlation. We display 16 universal chromatin states that show significant enrichment/depletion at *P*< 0.001 in any of the EWAS.

