## Supplementary Methods. Software for Xenopus clocks. for "DNA methylation clocks for clawed frogs reveal evolutionary conservation of epigenetic ageing"

Here we present details on the construction and software for our epigenetic clocks for Xenopus

A) all Xenopus tissues (both Laevis and Tropicalis combined),
B) all tissues from X. laevis,
C) all tissues from X. tropicalis,
D) all tissues from young Xenopus (age <= 2 years).
E) human-Xenopus clocks for chronological age: version 1 and version 2
F) human Xenopus clocks for relative age: version 1 and version 2

### Technical details surrounding epigenetic clocks

#### Statistical methods used for building the clocks

Since a mean methylation level of 0.5 is usually associated with a non detectable CpG (i.e. a CpG whose sequence does not map to the species genome), we restricted the analysis to cytosines whose mean methylation (across all Xenopous tissues) was outside of [0.4,0.6].

The epigenetic clocks were used by employing a single elastic net regression model analysis (R function glmnet). We use used Leave-one-out analysis (LOO) using a single lambda value. We chose the following parameters for the glmnet R function (Alpha: 0.5, CV Fold: 10, Lambda choice for Clock: 1 standard error above minimum CV-MSE).

#### Covariates and coefficient values of the frog clocks

The coefficient values of the clocks are specified in **Supplementary Table**.

1) The Xenopus clock (both tissues from Laevis and Tropicalis combined) is based on 58 CpGs whose coefficient values are specified in the column " XenopusClockLogLinear". Age transformation=log linear described below.
2) The Laevis (African clawed frog) clock (all tissues from Laevis) is based on 39 CpGs whose coefficient values are specified in the column "LaevisAfricanFrog". No age transformation.
3) The Tropicalis (Western clawed frog) clock (all tissues from Tropicalis) is based on 29 CpGs whose coefficient values are specified in the column "TropicalisWesternFrog". No age transformation.
4) The Young Xenopus clock (both tissues from Laevis and Tropicalis combined) is based on 33 CpGs whose coefficient values are specified in the column "YoungXenopusClockLogLinear". Age transformation=log linear described below.
5) The human Xenopus clock for chronological age is based on 228 CpGs whose coefficient values are specified in the column " HumanXenopusLogLinearVersion1". Age transformation=log-linear described below. Version 1 of this clock was used in the main text of the article by Zoller et al.
6) The human Xenopus clock for relative age is based on 213 CpGs whose coefficient values are specified in the column " HumanXenopusRelativeAgeVersion1". RelativeAge=Age/maxLifespan where the maximum lifespan of humans was set to 122.5, max lifespan of Laevis was set to 30.3 years, and max lifespan of Tropicalis=16 years. The estimates of maximum lifespan should be interpreted as mathematical parameters of the regression model.

Version 1 of this clock was used in the main text of the article by Zoller et al.

7) Version 2 of the human Xenopus clock for chronological age is based on 274 CpGs whose coefficient values are specified in the column " HumanXenopusLogLinearVersion2". Age transformation=log-linear described below. This clock has not yet been described. We have added it as "bonus material" for the Xenopus aficionado.
8) Version 2 of the human Xenopus clock for relative age is based on 500 CpGs whose coefficient values are specified in the column " HumanXenopusRelativeAgeVersion2". This clock has not yet been described. We have added it as "bonus material" for the Xenopus aficionado.

#### General description of age transformation

The human-frog clocks for chronological age used log linear transformations that are similar to those employed for the HUMAN pan tissue (Horvath 2013) (27).

An elastic net regression model (implemented in the glmnet R function) was used to regress a transformed version of age on the beta values in the training data. The glmnet function requires the user to specify two parameters (alpha and beta). Since I used an elastic net predictor, alpha was set to 0.5. But the lambda value of was chosen by applying a 10-fold cross validation to the training data (via the R function cv.glmnet).

The elastic net regression results in a linear regression model whose coefficients b_0_, b_1_, . . . , relate to transformed age as follows
*F*(chronological age)=*b*_0_*+b*_1_*CpG*_1_*+ . . . +b*_p_*CpG*_p_+error

Note that the intercept term is denoted by b_0_. The coefficient values can be found in Supplementary Table. Based on the coefficient values from the regression model, DNAmAge is estimated as follows
*DNAm*Age=$F^{-1}$(*b*_0_*+b*_1_*CpG*_1_*+ . . . +b*_p_*CpG*_p_),

where $F^{-1}\left( y \right)$ denotes the mathematical inverse of the function F(.). Thus, the regression model can be used to predict to transformed age value by simply plugging the beta values of the selected CpGs into the formula.

#### Defining Properties of the log linear transformation

As indicated by its name, the “log-linear” function, has a logarithmic dependence on age before the average age of sexual maturity (of the species) and a linear dependence after Age at Sexual Maturity (of the species). For the human-frog clocks we used the following averages at sexual maturity (in units of years): 13.5 years for humans, 1.0 years for Laevis, and 0.375 for Tropicalis.

We used a piecewise transformation, parameterized by Age of Sexual Maturity ($A$).

The transformation is F(x), given by

$$F\left( x \right)=g\left( \frac{x+1.5}{A+1.5} \right)\text{ where }g\left( t \right)= \left\{ \begin{aligned} \begin{aligned} \begin{aligned} \log\left( t \right), for 0\leq t\leq1 \\ t-1, for 1\leq t \end{aligned} \end{aligned} \end{aligned} \right.$$

Explicitly, F(x) is given by

$$F\left( x \right)=\left\{ \begin{aligned} \begin{aligned} \begin{aligned} \log\left( \frac{x+1.5}{A+1.5} \right), for 0\leq x\leq A \\ \frac{x-A}{A+1.5}, for A\leq x \end{aligned} \end{aligned} \end{aligned} \right.$$

In order to use this transformation to predict Age on *new samples*, one needs to use the *inverse* transformation, F^-1^(y), given by

$$F^{-1}\left( y \right)= \left\{ \begin{aligned} \begin{aligned} \begin{aligned} \left( A+1.5 \right)*\text{exp}\left( y \right)-1.5, for y\leq0 \\ (A+1.5)y+A, for y\geq0 \end{aligned} \end{aligned} \end{aligned} \right.$$

For predicting age, apply the inverse transformation to coefficient-weighted sum. That is,

$$DNAmAge=F^{-1}\left( x*\beta\right)$$

where $\beta$ is the vector of coefficients and $x$ is the vector of methylation values, with an intercept term.

#### R Implementation of the log linear transformation

```
### Applies the log linear transformation to the input vector x, i.e. to Age
F= Vectorize(function(x, maturity, ...) {
  if (is.na(x) | is.na(maturity)) {return(NA)}
  k <- 1.5
  y <- 0
  if (x < maturity) {y = log((x+k)/(maturity+k))}
  else {y = (x-maturity)/(maturity+k)}
  return(y)
})
### Inverse log linear transformation
F.inverse= Vectorize(function(y, maturity, ...) {
  if (is.na(y) | is.na(maturity)) {return(NA)}

  k <- 1.5
  x <- 0
  if (y < 0) {x = (maturity+k)*exp(y)-k}
  else {x = (maturity+k)*y+maturity}
  return(x)
})
```

#### The DNAm Age estimate is estimated in two steps.

First, one forms a weighted linear combination of the CpGs whose details can be found in Supplementary Table 1.

The table reports the probe identifier (cg number) used in the custom Infinium array (HorvathMammalMethylChip40). The weights used in this linear combination are specified in the respective column entitled "Coef.".

The formula assumes that the DNA methylation data measure "beta" values but the formula could be adapted to other ways of generating DNA methylation data.

**Pseudo R code**

```
# R function for multivariate regression model
multivariatePredictorCoef=function(dat0, datCOEF,imputeValues=FALSE) {
datout=data.frame(matrix(NA,nrow=dim(dat0)[[2]]-1,ncol=dim(datCOEF)[[2]]-1 ))
match1=match(datCOEF[-1,1],dat0[,1] )
if (   sum(!is.na(match1))==0 ) stop("Input error. The first column of dat0 does not contain CpG identifiers (cg numbers).")
dat1=dat0[match1,]
row.names1=as.character(dat1[,1])
dat1=dat1[,-1]
if (imputeValues ){dat1=impute.knn(data=as.matrix(dat1) ,k = 10)[[1]]}
for (i in 1:dim(dat1)[[2]] ){ for (j in 2:dim(as.matrix(datCOEF))[[2]] ){
datout[i,j-1]=sum(dat1[,i]* datCOEF[-1,j],na.rm=TRUE)+ datCOEF[1,j]}}
colnames(datout)=colnames(datCOEF)[-1]
rownames(datout)=colnames(dat0)[-1]
datout=data.frame(SampleID= colnames(dat0)[-1],datout)
datout
} # end of function

# read in supplementary table
datCoef=read.csv("TableS.csv")

The first columns should read as follows
names(datCoef)
  [1] "var"
  [2] "Coef.XenopusClockLogLinear"
  [3] "Coef.LaevisAfricanFrog"
  [4] "Coef.TropicalisWesternFrog"
  [5] "Coef.YoungXenopusClockLogLinear"
  [6] "Coef.HumanXenopusLogLinearVersion1"
  [7] "Coef.HumanXenopusRelativeAgeVersion1"
  [8] "Coef.HumanXenopusLogLinearVersion2"
  [9] "Coef.HumanXenopusRelativeAgeVersion2"

# Restrict attention to the first 5 columns
datCoef=datCoef[,c(1:9)]

# assume the first column of dat0 contains the CpG identifiers
match1=match(datCoef[-1,1],dat0[,1] )
missingProbes= as.character(datCoef[-1,1] )[is.na(match1)]

dat1=dat0[match1,]
# data frame with predicted values.
datPredictions=multivariatePredictorCoef(dat1,datCOEF=datCoef,imputeValues=FALSE)

#let's relabel the columns by replacing "Coef" with "DNAm" since the columns contain estimates of age or relative age instead of coefficient values

colnames(datPredictions)=gsub(pattern="Coef", replacement="DNAm", x=colnames(datPredictions))

# We need to transform the human frog clock for chronological age using the inverse of the log linear #transformation.

For LAEVIS, the age at sexual maturity has to be set to 1.0 years.
datPredictions$DNAm.HumanXenopusLogLinearVersion1= F.inverse(datPredictions$DNAm.HumanXenopusLogLinearVersion1, maturity=1.0)
datPredictions$DNAm.XenopusClockLogLinear= F.inverse(datPredictions$DNAm.DNAm.XenopusClockLogLinear, maturity=1.0)
datPredictions$DNAm.YoungXenopusClockLogLinear= F.inverse(datPredictions$DNAm.DNAm.YoungXenopusClockLogLinear, maturity=1.0)

For Tropicalis, the age at sexual maturity has to be set to 0.375
datPredictions$DNAm.YoungXenopusClockLogLinear= F.inverse(datPredictions$DNAm.DNAm.YoungXenopusClockLogLinear, maturity=0.375)
datPredictions$DNAm.HumanXenopusLogLinearVersion1= F.inverse(datPredictions$DNAm.HumanXenopusLogLinearVersion1, maturity= 0.375)
datPredictions$DNAm.XenopusClockLogLinear= F.inverse(datPredictions$DNAm.DNAm.XenopusClockLogLinear, maturity=0.375)


#The data frame "datPredictions" contains the age estimates in units of years and relative age estimates.

Legal boiler plate: No warranties of any kind.
```
